# Low concentrations of the food contaminant Deoxynivalenol trigger apoptosis and impair GnRH-induced LH secretion in pituitary gonadotrope-like cells

**DOI:** 10.1101/2024.08.13.607800

**Authors:** Guodong Cai, Lingchen Yang, Francis Marien-Bourgeois, Derek Boerboom, Gustavo Zamberlam, Imourana Alassane-Kpembi

**Affiliations:** College of Veterinary Medicine, Yangzhou University, Yangzhou 225009, Jiangsu, China; College of Veterinary Medicine, Hunan Agricultural University, Changsha 410128, Hunan, China; Centre de Recherche en Reproduction et Fertilité (CRRF), Département de Biomédecine vétérinaire, Faculté de Médecine vétérinaire, Université de Montréal, Saint-Hyacinthe H3T 1J4, Québec, Canada

**Keywords:** Mycotoxins, deoxynivalenol, pituitary gland, LβT2 cells, MAPK signaling pathway, gonadotropins, LH

## Abstract

The *Fusarium* mycotoxin deoxynivalenol (DON) is one of the most frequently occurring food contaminants. Nearly all individuals are exposed to DON, due to it widespread presence in grains and grain-based products. Chronic DON poisoning is associated with growth retardation, immunotoxicity as well as impaired reproduction and fetal development. At the molecular level, DON alters intracellular signaling by activating mitogen-activated protein kinases (MAPKs) that modulate cell growth, differentiation, and apoptosis. Of note, these MAPKs are also critical mediators of gonadotrophin-releasing hormone (GnRH)-induced synthesis and secretion of follicle-stimulating hormone (FSH) and luteinizing hormone (LH) by pituitary gonadotrope cells. So far, no research has explored the potential endocrine-disrupting effects of DON on pituitary gonadotropins production. Herein, we show the first evidence that DON affects LH production by the immortalized gonadotrope-like cell line LβT2 in a concentration-dependent manner. Taken together, our experiments demonstrated that low concentrations of DON affect GnRH-induced signaling through a mechanism that, at least in part, involves apoptosis and inhibition of GnRH-induced phosphorylation of ERK-MAPK. Consequently, DON also affects the GnRH-induced expression of *Cga* and *Lhb*, two critical genes for LH synthesis and secretion by gonadotrope cells in mammals. This research broadens our knowledge of the toxicity of DON and brings a new depth to the potential neuroendocrine implications for reproduction.

**Graphical abstract:** 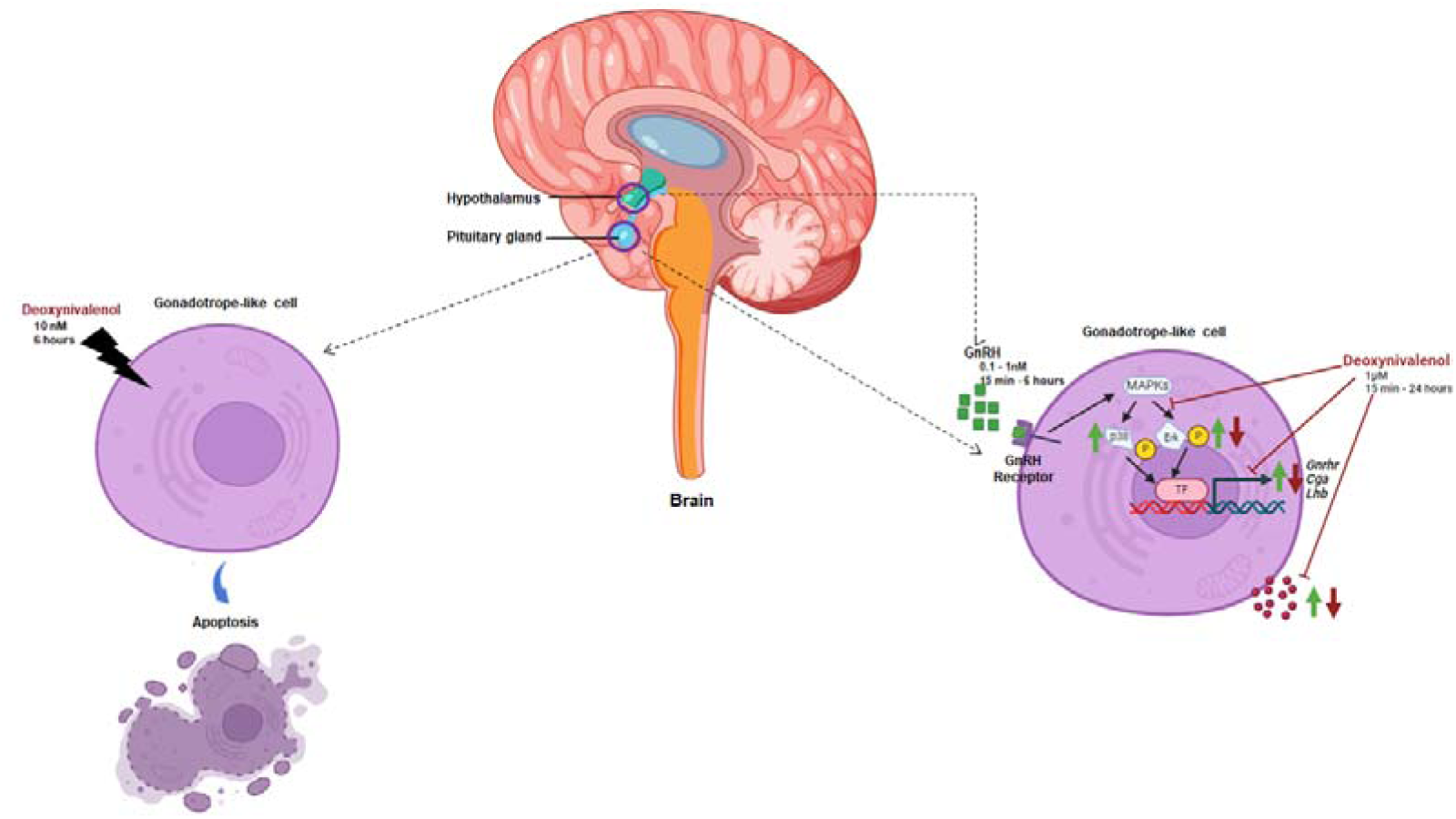

## 1. Introduction

Mycotoxins, as pervasive contaminants of foodstuffs, pose a substantial threat to both food security and human health. Among them, Deoxynivalenol (DON), produced by *Fusarium* fungi species, is one of the most prevalent, and represents a global health concern (Streit et al. 2012). Nearly all individuals worldwide are exposed to DON, and the current exposure levels necessitate taking vigilant steps to guarantee food safety (Mishra et al. 2020). Acute exposure to DON in experimental animal models cause symptoms such as diarrhea, vomiting, leukocytosis, hemorrhage, and even death, while chronic low-dose exposure is associated with anorexia, growth retardation, immunotoxicity and altered neuroendocrine signaling (Pestka 2010).

A negative impact of DON on reproductive functions primarily related to altered gonadal function has also been reported in several species (Pizzo et al. 2015; Sprando et al. 1999; Tassis et al. 2022). Interestingly, male rats exposed to DON presented altered serum levels of the gonadotropins follicle-stimulating hormone (FSH) and luteinizing hormone (LH), suggesting that DON could act directly at the level of the pituitary gland (Sprando et al. 2005).

At the cellular level, the primary mechanism of DON toxicity involves inhibiting protein and nucleic acid synthesis by binding to the ribosome and activating cellular kinases, including the mitogen-activated protein kinases (MAPKs) p38, c-Jun N-terminal kinase, and extracellular-signal-regulated kinase (ERK) (Payros et al. 2016). This mechanism influences various signal transduction pathways, including those related to cell proliferation, stress, inflammation, differentiation and apoptosis (Murphy and Blenis 2006). Most importantly, MAPKs are also critical mediators of GnRH-induced gonadotropin synthesis and secretion by pituitary gonadotrope cells (Kahnamouyi et al. 2018). Gonadotropins play a pivotal role in both male and female reproduction. In males, LH regulates the synthesis of androgens by the Leydig cells, whereas FSH promotes Sertoli cell function and thereby influences spermatogenesis (Ulloa-Aguirre and Lira-Albarrán 2016). In females, FSH is the key hormone for follicle growth, selection of the dominant follicle and oocyte maturation, while a midcycle surge of LH is required for oocyte meiotic maturation, successful ovulation, and luteinization of the follicle (Singh et al. 2023). LH and FSH are dimeric glycoproteins composed of a common alpha subunit (CGα) and distinct β subunits (LHβ and FSHβ), which confer biological specificity (Bernard and Tran 2013). The main determinant of gonadotropin synthesis is the level of expression of *LHB* and *FSHB.* The transcription of *LHB* is regulated mainly by hypothalamic gonadotropin-releasing hormone (GnRH) (Tremblay and Drouin 1999), which binds to its receptor (GnRHr) to activate the extracellular regulated kinases 1 and 2 (ERK1/ 2) via phosphorylation (Bliss et al. 2009; Liu et al. 2002). Yet, to the best of our knowledge, no research has explored the potential endocrine-disrupting effects of DON on pituitary gonadotropin production.

To start addressing this gap, our study investigated the effects of DON on pituitary gonadotrope cell viability, and its impact on GnRH-stimulated ERK signaling and LH secretion. As gonadotropes are a rare cell population in the pituitary gland (making their enrichment and primary culture considerable technical challenges), we employed the immortalized gonadotrope-like cell line LβT2. This cell line is derived from a mouse pituitary tumor, expresses GnRHr, CGα, LHβ and FSHβ and secretes LH and FSH upon stimulation (Graham et al. 1999; Turgeon et al. 1996). Therefore, this cell line serves as a crucial tool for the study of gonadotrope physiology, as well as for the identification of potential endocrine disruptors of gonadotropin production in mammals.

## 2. Materials and methods

### 2.1 Chemicals and Reagents

DON and dimethyl sulfoxide (DMSO) were obtained from Millipore Sigma Canada Ltd. (Ontario, Canada). GnRH was from Peptide Pros (Miami, USA). Dulbecco’s Modified Eagle’s Medium (DMEM), Fetal Bovine Serum (FBS), gentamycin and Trypsin were from Wisent (St-Bruno, Quebec, Canada).

### 2.2 Cell culture

After thawing, LβT2 cells were seeded in flasks and cultured in DMEM medium (Wisent, St-Bruno, Quebec, Canada) supplemented with 10% FBS and 4 µg/mL gentamycin and incubated at 37L with 5% CO_2_. The culture medium was refreshed every two days after cell adhesion to the flask was achieved. When the cells reached 80% confluence, they were trypsinized and seeded in a specific plate at a specific density, according to the needs of each experiment, as described below.

### 2.3 Cytotoxicity assessment

LβT2 cells were distributed in 96-well plates at 5×10^3^ cells in 100 μL culture medium per well and allowed to attach. The following day, the culture medium was aspirated, and the cells were exposed to 100 μL of new medium with graded concentrations of DON: 0 nM (control), 1 nM, 10 nM, 10^2^ nM, 10^3^ nM and 10^4^ nM for 24 or 48h. Cell viability was then determined by CellTiter Blue (Promega, Madison, WI) as previously described by Cai et *al*. (2023).

### 2.4 Real-time apoptosis and necrosis assay

The Real Time-Glo™ Annexin V Apoptosis and Necrosis Assay (Promega, Madison, WI, Cat#: JA1011) was used to simultaneously analyze apoptosis and necrosis. This assay probes the presence of phosphatidylserine on the outer leaflet of the cell membrane during apoptosis, while producing a fluorescent signal upon the loss of membrane integrity because of the binding of the cell-impermeant pro-fluorescent DNA dye it contains.

For this experiment, LβT2 cells were seeded at 3×10^4^ cells per well in 100 μL culture medium in white clear-bottom 96-well plates (Costar, Cambridge, MA, USA) and incubated in an atmosphere of 5% CO2 at 37 °C. After 48 h, 50 μL of the existing cell medium was carefully withdrawn and replaced by an equivalent volume of fresh medium without (control group) or with 40 nM DON. In each well, 100 μL of a 2X reagent solution was dispensed, according to the manufacturer’s instructions, resulting in a final liquid volume of 200 μL per well. This ensured a final mycotoxin concentration of 10 nM in the treated well. Subsequently, the plates were placed in a 37°C incubator with 5% CO2. Measurements were taken at 2 h, 6 h and 24 h post-treatment, to assess membrane integrity through fluorescence (excitation/emission wavelength: 485 ± 10 nm / 525 ± 10 nm) and phosphatidylserine translocation through the luminescent signal generated by the Annexin V-dependent assembly of two luciferase fragments.

### 2.5 LβT2 cell treatments for MAPK activation

LβT2 cells were distributed in 24-well plates at 1.5×10^5^ cells in 300 μL of medium per well and cultured for 24 hours. Then the cells were either directly stimulated with graded concentrations of GnRH from 0.001nM to 0.1nM for 15 min; or pre-treated with DON at concentrations of 1 μM or 10 μM for 15 min before stimulation with the highest concentration of GnRH (0.1nM) for 15 min. Following GnRH challenge, cells were collected, and cellular proteins were extracted using a radioimmunoprecipitation assay (RIPA) buffer. The protein concentrations of the samples were quantified using a bicinchoninic acid (BCA) protein quantification kit from Thermo Fisher Scientific (Waltham, Massachusetts, USA). Twenty μg of protein from each sample were separated using a Precast Midi Protein Gel (BIO-RAD, Hercules, California, USA) and then transferred to polyvinylidene fluoride (PVDF) membranes (BIO-RAD, Hercules, California, USA). These were probed using primary antibodies specific to total ERK and p38, phosphorylated ERK and p38, and β-tubulin. Detection was done using a peroxidase-conjugated secondary antibody (Jackson ImmunoResearch Inc, USA), and the antigen-antibody complexes were visualized using an enhanced chemiluminescence assay (ECL, Invitrogen, Carlsbad, California, USA).

### 2.5 LβT2 cell treatments for gene expression and hormonal assay

To determine effects of DON on the expression pattern of gonadotropin synthesis-related genes and LH secretion, LβT2 cells were seeded and allowed to attach in 6-well plates at 1×10^6^ cells per well. On the next day, the cell supernatant was carefully aspirated and replaced with new culture medium with graded concentrations of DON ranging from 1 nM to 10 µM for 24 h. Following this pre-treatment with DON, the cells were stimulated with 1nM GnRH for 6 h. While in one experiment cells were harvested for total RNA extraction, in another experiment medium was collected for LH measurement.

#### 2.5.1 Total RNA extraction and real-time PCR

The expression of the gene encoding the GnRH receptor (*Gnrhr*) and the gonadotropin-related genes *Cga*, *Lhb*, and *Fshb* was analyzed. For this, total RNA was first extracted using Trizol as per the guidelines provided by the manufacturer (BIO-RAD). Following purity and concentration assessment on a Nanodrop spectrophotometer, total RNA (1µg) was reverse transcribed into cDNA, using the Advanced cDNA synthesis Kit (Wisent Bioproducts, St-Bruno, QC, Canada) according to the manufacturer’s instructions. RT-qPCR was run on a CFX96 Touch Real-Time PCR Detection System (BIO-RAD) using qPCR Mastermix with Supergreen (Wisent Bioproducts, St-Bruno, QC, Canada) following the manufacturer’s protocol. To ensure specificity of PCR products, melting curves were analyzed and the amplicons were sequenced to verify authenticity. Experimental samples were run in duplicate and data were expressed relative to *Rpl19* as housekeeping gene, and normalized to a calibrator sample using the Pfaffl method with correction for amplification efficiency (Pfaffl 2001). Primer details for both housekeeping and target genes are provided in Table 1.

**Table 1.**
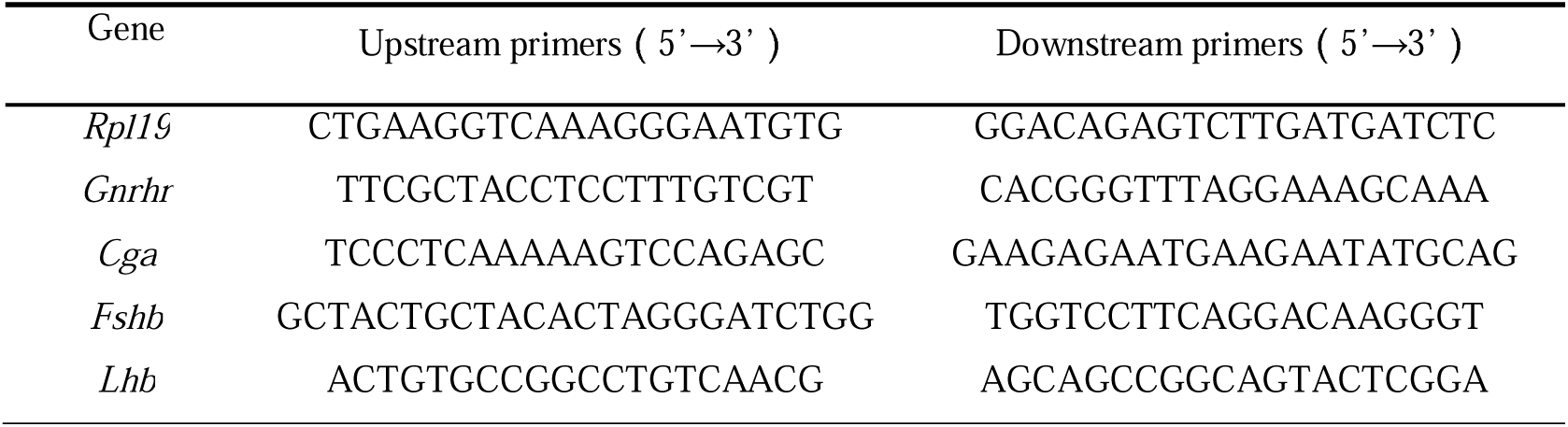
Primer sequences of RT-qPCR.

#### 2.5.2 LH measurement

LH levels in cell culture medium samples were evaluated using the Ultra-Sensitive Mouse and Rat LH ELISA performed by the Center for Research in Reproduction at the Ligand Assay and Analysis Core Laboratory of the University of Virginia. The capture monoclonal antibody (anti-bovine LH beta subunit, 518B7; RRID:AB_2784498) was provided by Janet Roser, University of California. The detection polyclonal antibody (rabbit LH antiserum, AFP240580Rb; RRID:AB_2784499) was provided by the National Hormone and Peptide Program (NHPP). HRP-conjugated polyclonal antibody (goat anti-rabbit) was purchased from DakoCytomation (Glostrup, Denmark; D048701-2; RRID:AB_2784500). Mouse LH reference prep (AFP5306A; NHPP) was used as the assay standard. The limit of quantitation was 0.016 ng/ml.

### 2.6 Statistical analysis

All statistical analyses were performed with Prism software 8.0.2 for Windows (GraphPad). The reported values are the means ± standard error of the mean (SEM) of at least three independent experiments, each with triplicate wells per experimental condition. Data were transformed to logarithms if they were not normally distributed (Shapiro–Wilk test). Two-tailed *t* tests or one-way ANOVAs followed by Tukey’s multiple-comparison tests were run for comparisons between the experimental conditions. Differences with a *p* value < 0.05 were considered statistically significant.

## 3 Results

### 3.1 Low concentrations of DON alter gonadotrope-like cell viability

The cytotoxicity of DON for LβT2 cells was assessed using the CellTiter-Blue® Cell Viability Assay. The results indicated decreased viability of LβT2 cells following exposure to increased graded concentrations of DON (Fig. 1). Interestingly, it was observed that viability was significantly different from control after 24 h (*p*<0.05) or 48 h (*p*<0.01) of exposure even the lowest tested concentration of 1 nM. The decline in cell viability showed a clear concentration-dependent pattern at both 24 and 48 h, reaching its nadir at 10^4^ nM DON for both exposure durations (*p*<0.01). Furthermore, the results also indicated that with longer exposure time (48h *vs.* 24h), the impact on LβT2 cell viability was more profound. This shows a concentration and time-dependent reduction in gonadotrope-like cell viability in response to DON starting from the nanomolar range.

**Figure 1.**
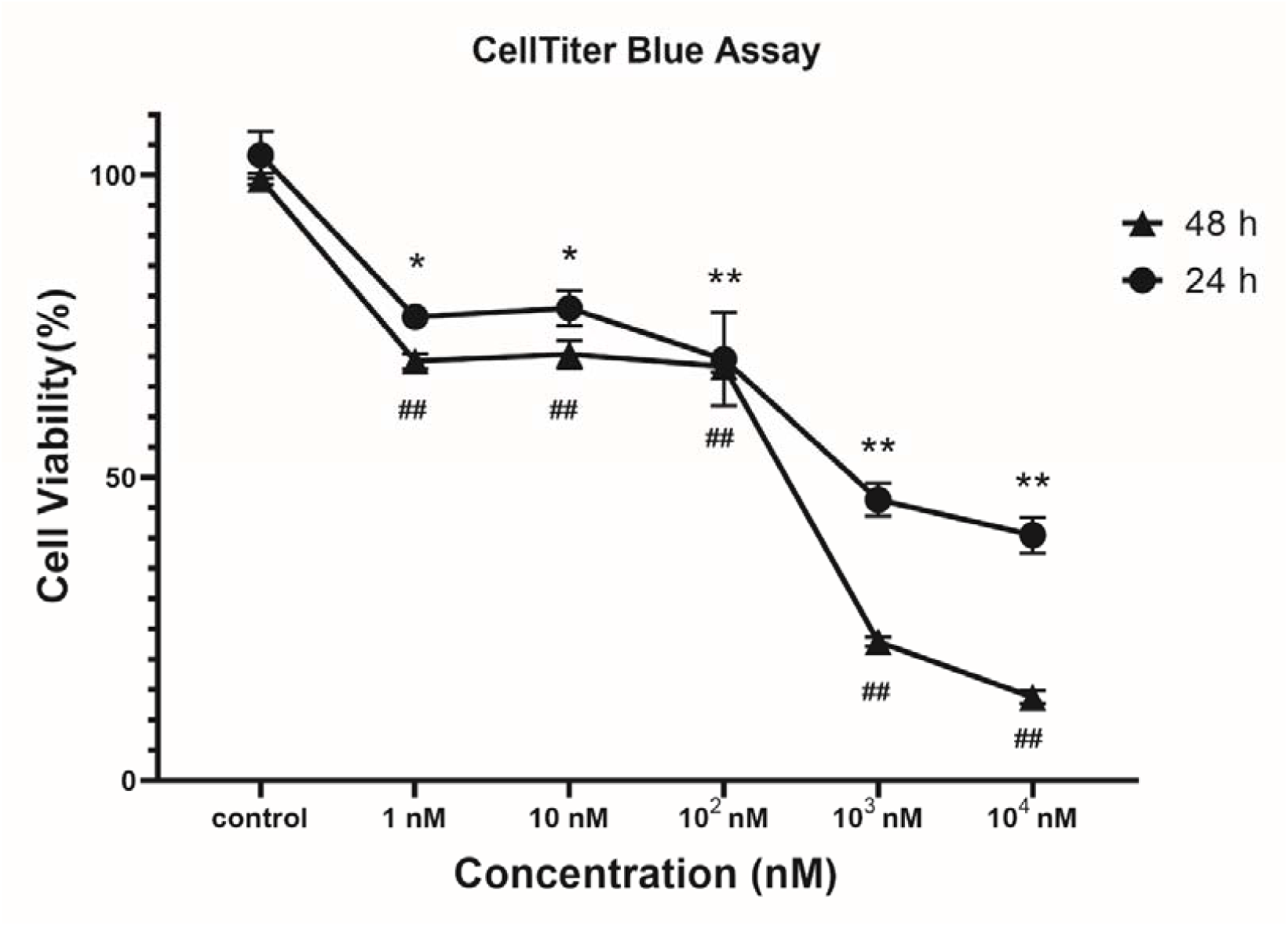
Cytotoxic effects of DON on LβT2 cells. Cell viability (%) of was determined by CellTiter Blue in LβT2 cell exposed to DON for 24 and 48 h in triplicate, and the experiment was repeated 3 times (n=3). Significant differences post 24 h treatments compared with control, \**p < 0.05*; ** *p < 0.01*. Significant difference post 48 h treatments compared with control, **^#^***p < 0.05*; **^##^** *p < 0.01*.

### 3.2 DON induces apoptosis in gonadotrope-like cells

To further elucidate the mechanism of DON-induced cytotoxicity on LβT2 gonadotrope-like cells, we then conducted a real-time assay with the objective of determining the temporal dynamics of translocation of phosphatidylserine and loss of membrane integrity following exposure to this mycotoxin. Translocation of phosphatidylserine to the outer leaflet of the plasma membrane was revealed through annexin V binding coupled with a luminescent signal emission (Fig. 2a), while loss of membrane integrity was associated with a fluorescent signal resulting from DNA staining by a cell-impermeant reagent (Fig. 2b). Apoptotic cell death is therefore characterized by a temporal lag between phosphatidylserine translocation and the compromise of membrane integrity, whereas necrotic cell death is characterized by simultaneous annexin V binding and the disruption of membrane integrity.

**Figure 2.**
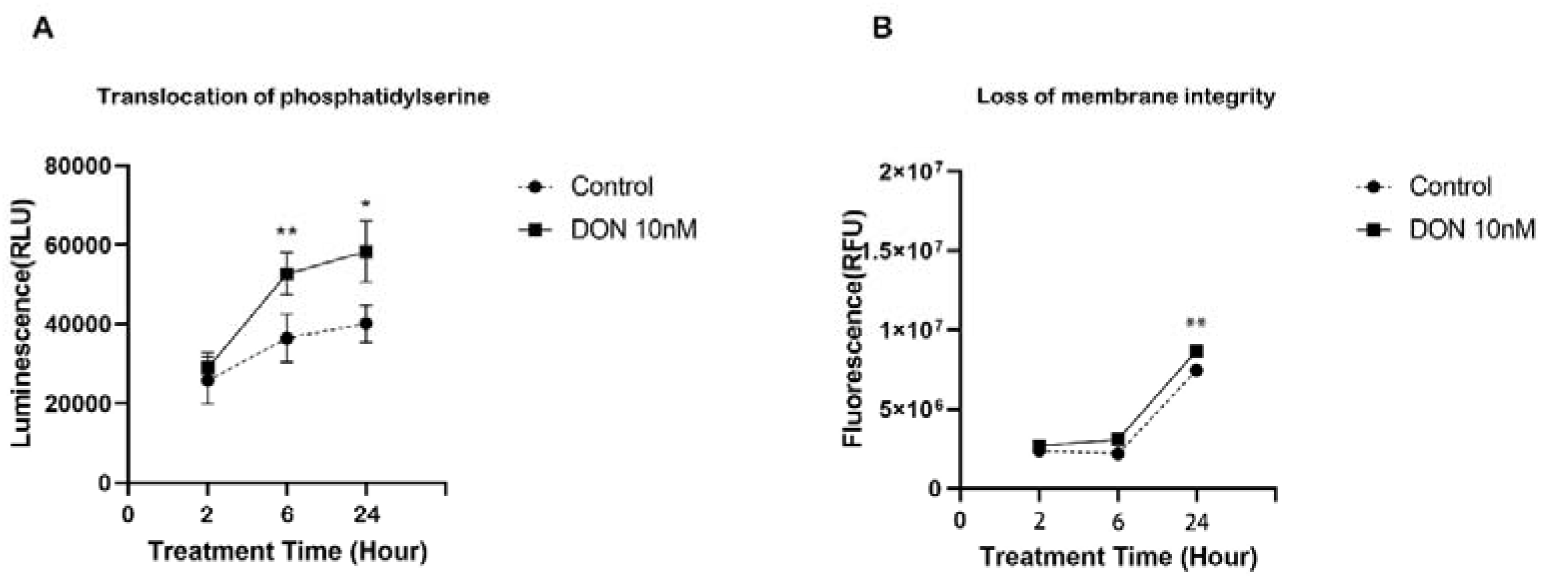
DON induces apoptosis in LβT2 cells. RealTime-Glo™ Annexin V Apoptosis and Necrosis Assay detection of apoptosis (A) and secondary necrosis (B) in LβT2 cells treated with DON (10 nM) in triplicate, and the experiment was repeated 3 times (n=3). Significant differences post 24 h treatment compared with control, \**p < 0.05*; ** *p < 0.01*.

For this experiment, we chose 10 nM of DON in the treated group. According to the previous experiment, at this dose, DON already alters cells viability at both 24 and 48h post-exposure. At the 6-h exposure time point, it was already possible to observe a significant increased phosphatidylserine translocation compared to the control (*p*<0.01), while no significant difference in membrane integrity loss was observed (*p*>0.05). However, both phosphatidylserine translocation and membrane integrity compromise showed statistically significant differences between the DON-treated cells and the control at the 24-h exposure. These findings indicate that exposure to 10 nM DON for 6 h induced apoptosis without triggering necrosis.

### 3.3 DON attenuates GnRH-induced phosphorylation of Erk but not p38 in gonadotrope-like cells

To start exploring the potential negative impact of DON on pituitary gonadotropin production-related signaling, we investigated the effects of DON exposure on the activation of MAPKs by GnRH in gonadotrope-like cells. For this, LβT2 cells were challenged with distinct doses of GnRH ranging from 0.001nM to 0.1nM for 15 min. However, only GnRH at the dose of 0.1 nM significantly increased the phosphorylation levels for Erk (Fig. 3; *p*<0.01) and p38 (Fig. 4; *p*<0.01) as expected. Most interestingly, when LβT2 cells were pre-exposed to DON at concentrations of 1 μM and 10 μM before challenging with 0.1 nM GnRH, only the phospho-Erk (Fig. 3; *p*<0.05) but not the phospho-p38 (Fig. 3; *p*<0.01) increase was attenuated. Together, these results indicate that DON interferes with GnRH-stimulated ERK signaling.

**Figure 3.**
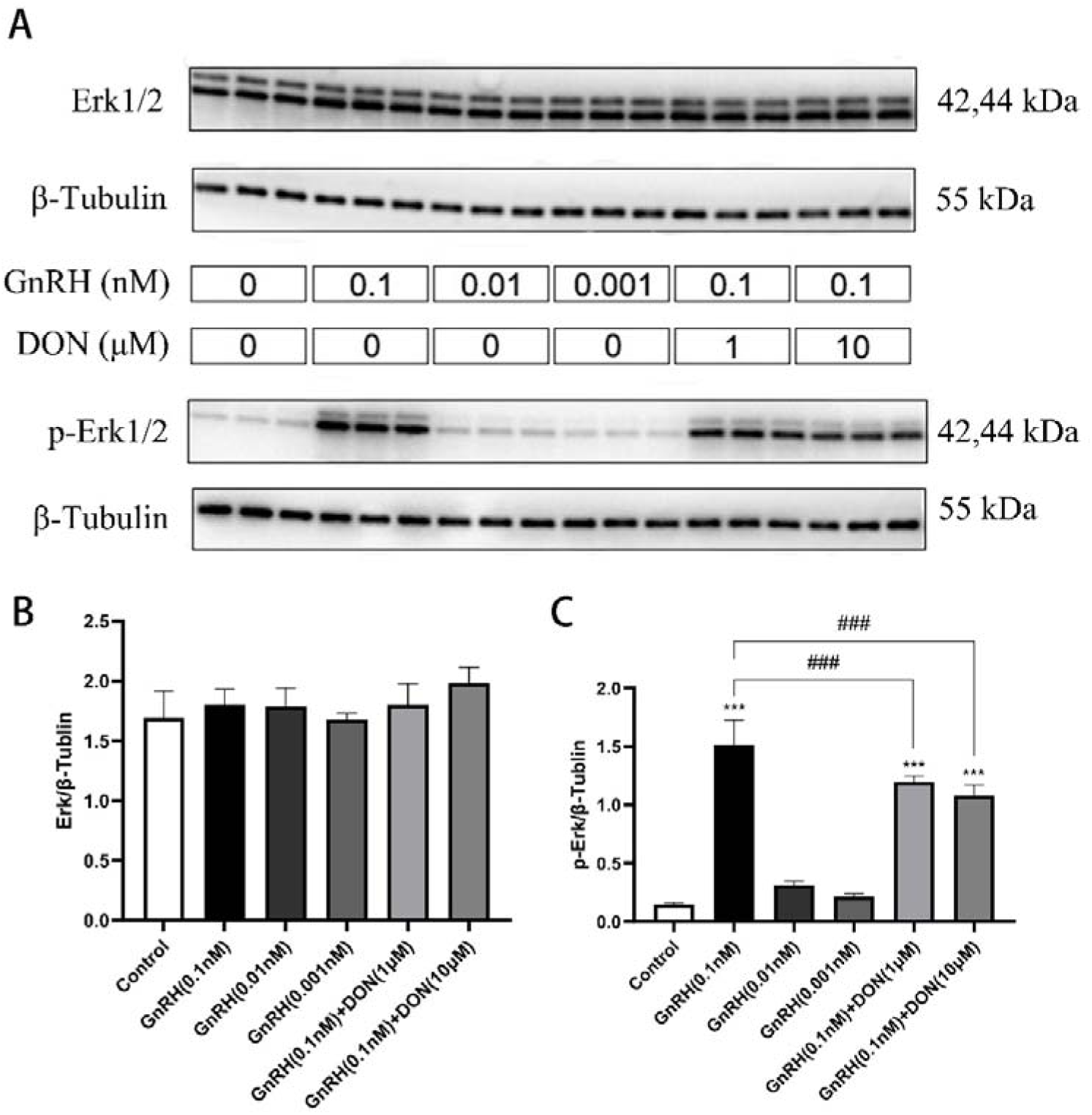
Effects of DON on the expression and phosphorylation of ERK 1/2 proteins in LβT2 cells stimulated by GnRH. LβT2 cells were pre-exposed for 24 h to DON at concentrations of 1 μM and 10 μM before challenging them with distinct doses of GnRH for 15 min. Protein levels were measured by Western blot and total and phosphorylated levels of ERK 1/2 were normalized to β-actin. The experiment was repeated 3 times (n=3). Significant effect of GnRH treatment compared with control, *** *p < 0.001*; Significant effect of DON treatment compared with GnRH alone, ^##^*p < 0.01*; ^###^ *p < 0.001*.

**Figure 4.**
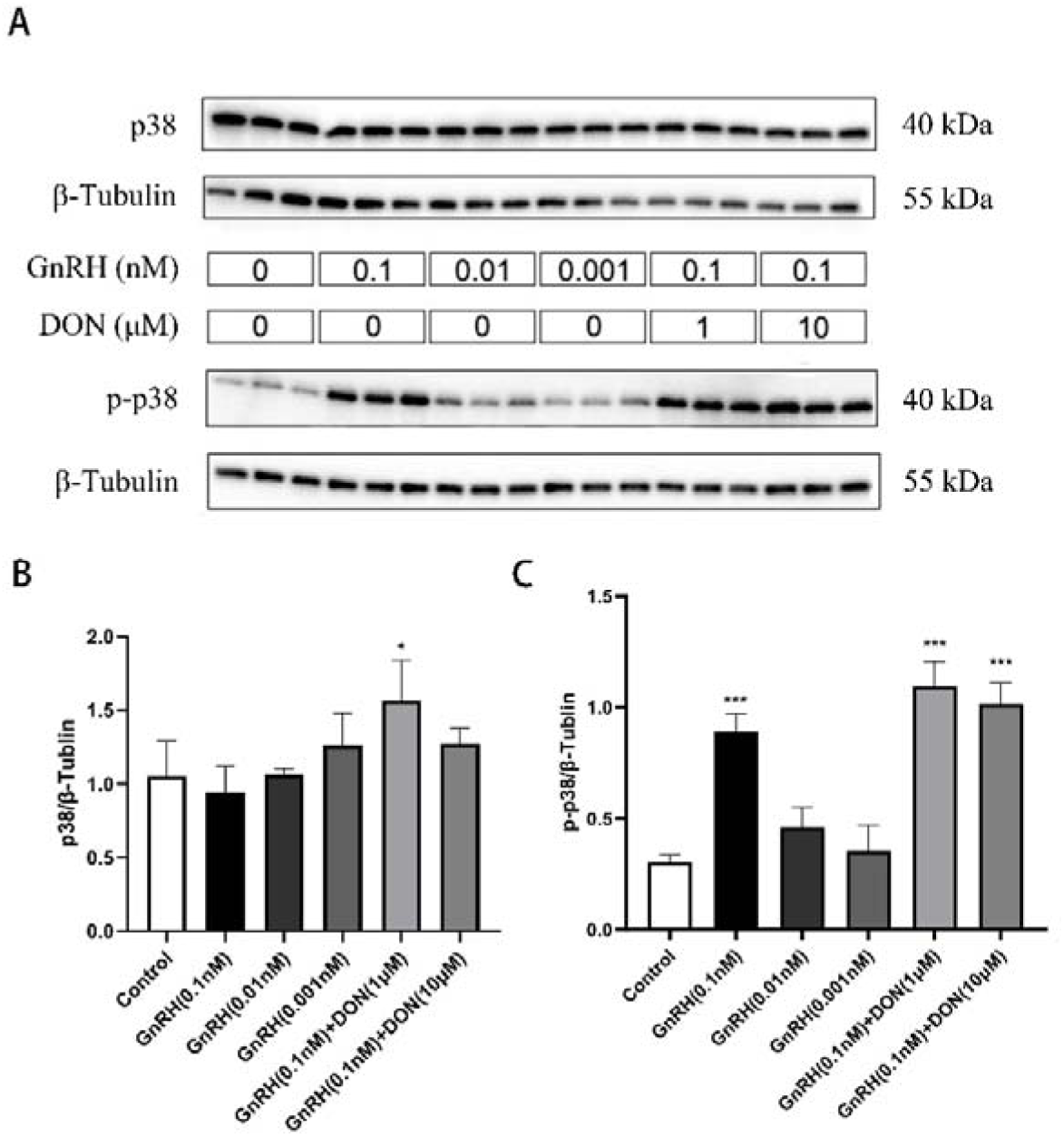
Effects of DON on the expression and phosphorylation of p38-MAPK protein in LβT2 cells stimulated by GnRH. LβT2 cells were pre-exposed for 24 h to DON at concentrations of 1 μM and 10 μM before challenging them with distinct doses of GnRH for 15 min. Protein levels were measured by Western blot and total and phosphorylated levels of p38-MAPK were normalized to β-actin. The experiment was repeated 3 times (n=3). Significant effect of GnRH and/or DON treatments compared with controls, \**p < 0.05*; *** *p < 0.001*.

### 3.4 DON attenuates the GnRH-induced expression of Cga and Lhb but not Gnrhr in gonadotrope-like cells

To better elucidate the impact of DON on GnRH-induced downstream signaling in gonadotrope-like cells, we also investigated the effects of DON on GnRH-induced expression of some of the genes considered essential for the synthesis of gonadotropins. For this, we first performed a preliminary experiment using the same doses of GnRH employed in the previous experiment. However, we did not observe a GnRH-induced increase of *Gnrhr, Cga* or *Lhb* mRNA levels at any dose (0.001 nM to 0.1 nM, data not shown). When we increased the GnRH dose to 1nM, the expression of these three genes was significantly augmented (Fig.5; *p*<0.05). Interestingly, while the GnRH-induced expression of *Cga* and *Lhb* was significantly attenuated by DON pre-treatment (*p*<0.05), *Gnrhr* was not (*p*>0.05). GnRH did not increase *Fshb* expression, which is a typical outcome when only a single pulse of GnRH is used (Lawson et al. 2007). We nonetheless were able to determine that DON does not affect the basal expression of either *Fshb* (*p*>0.05) nor of that of the other gonadotropin-synthesis related genes (*p*>0.05).

**Figure 5.**
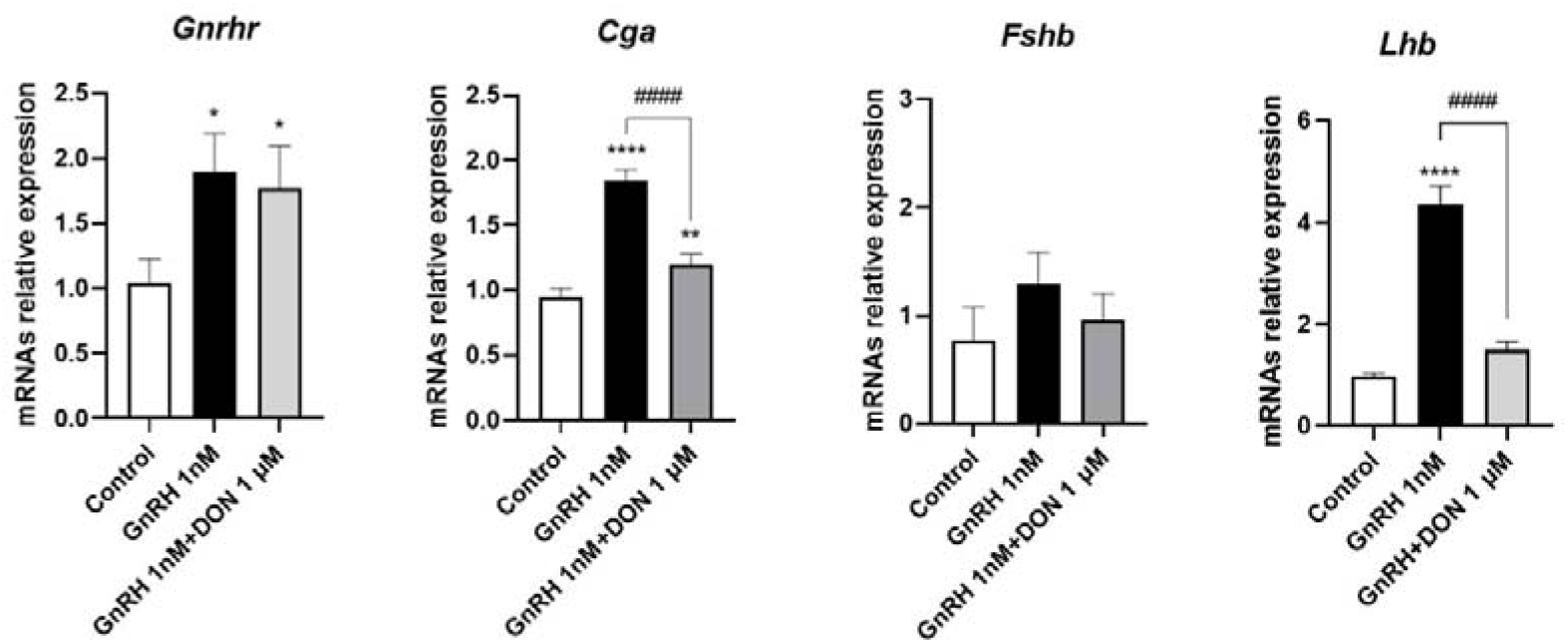
Effects of DON on *Gnrhr*, *Cga*, *Fshb* and *Lhb* gene expression in LβT2 cells stimulated by GnRH. LβT2 cells were pre-exposed for 24 h to DON at concentrations of 1 μM or 10 μM before challenging them with distinct doses of GnRH for 6 h. Cells were collected for RNA extraction and mRNA levels of *Gnrhr*, *Cga*, *Fshb* and *Lhb* were assessed by RT-qPCR and normalized to the housekeeping gene *Rpl19*. The experiment was repeated 3 times (n=3). Significant effects of GnRH treatment compared with untreated control, \**p < 0.05*; \*\**p < 0.01*; *** *p < 0.0001*. Significant effect of DON+GnRH treatment relative to GnRH-treated control, ^####^*p < 0.0001*.

### 3.5 LH secretion by gonadotrope-like cells is dose-dependently inhibited by DON

Together, all the previous experiments demonstrated that DON affects GnRH-induced signaling in gonadotrope-like cells trough a mechanism that, at least in part, involve apoptosis and attenuation of GnRH-induced Erk phosphorylation. Consequently, DON also affects the GnRH-induced expression of *Cga* and *Lhb*, two critical genes for LH synthesis and secretion by gonadotrope cells. To ultimately determine if DON would indeed affect LH secretion by gonadotropes, we then performed an experiment in which LβT2 cells were pre-exposed to graded concentrations of DON (ranging from 1 to 10^4^ nM) and then challenged with 1 nM GnRH (Fig. 6). As expected, GnRH at this concentration significantly induced LH release into the culture medium in comparison to control (*p*<0.05). Interestingly, at 1nM DON was not able to attenuate GnRH-induced LH production (*p*>0.05), but at 10^3^ nM and 10^4^ nM, DON inhibited it successfully (*p*<0.05). Together, these results clearly indicate the DON exerts a concentration-dependent negative effect on GnRH-induced LH secretion.

**Figure 6.**
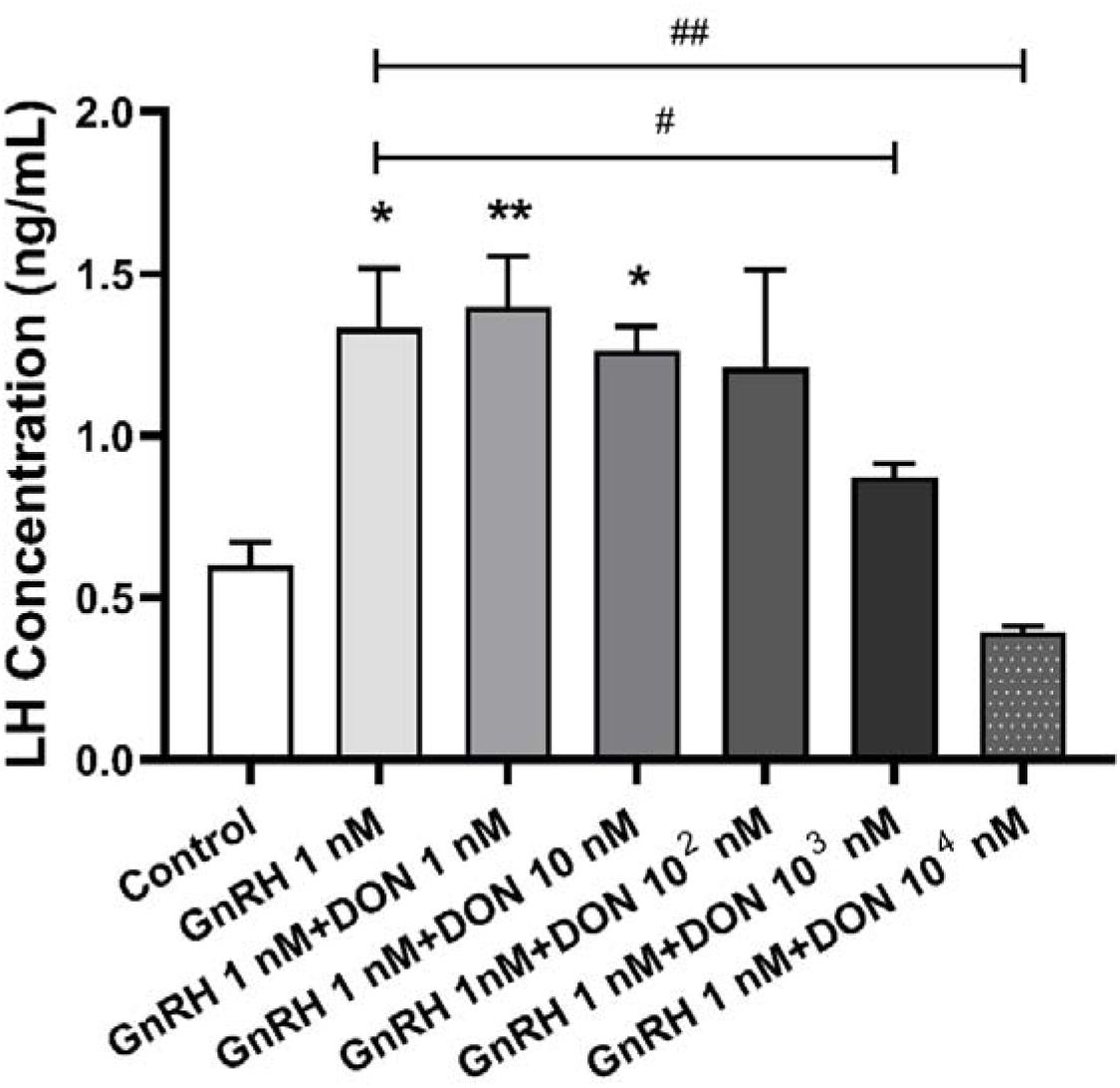
Effects of DON on LH concentration in culture medium of LβT2 cells stimulated by GnRH. LβT2 cells were treated or not with graded concentrations of DON for 24 hours, prior to stimulation with 1nM GnRH for 6 hours. Cell culture media samples were collected and then analyzed by Ultra-Sensitive Mouse and Rat LH ELISA. The experiment was repeated 3 times (n=3). Significant differences of GnRH and/or DON treatments compared with untreated control, \**p < 0.05*; ** *p < 0.01*. Significant effect of DON+GnRH treatment relative to GnRH-treated control, ^#^*p < 0.05*); ^##^ *p < 0.01*.

## Discussion

The present study provides strong evidence for the potential endocrine-disrupting effects of DON exposure on pituitary gonadotropin production in mammals. We demonstrate herein for the first time that exposure to realistic concentrations (i.e. low concentrations observed in the diet) of the type B trichothecene mycotoxin DON can not only be cytotoxic to gonadotrope-like cells, but can impair GnRH-induced LH secretion in this cell type in a concentration-dependent manner.

A growing body of literature is pointing to the impact of *Fusarium* mycotoxins on the hypothalamic–pituitary–gonadal (HPG) axis. The type A trichothecene mycotoxin T-2 was reported to induce alterations upstream in the rat HPG axis, through increase of hypothalamic GnRH synthesis, and downstream through precocious onset of the pubertal development of reproductive organs (Yang et al. 2016). Downstream of the HPG axis, the type B trichothecene DON has been shown to have a negative impact on bovine granulosa cell steroidogenesis and survival (Guerrero-Netro et al. 2015). In rats, DON also increased progenitor Leydig cell proliferation, but inhibited their maturation and function, including steroidogenesis (Yang et al. 2024). Only a few studies have investigated the toxicity of mycotoxins midstream of the HPG axis. DON inhibited the growth of rat pituitary GH3 cells (Liu et al. 2020). In porcine gonadotrope cells, the *Fusarium* toxins ZEA and α-ZOL specifically inhibited FSH synthesis and secretion, but not that of LH (He et al. 2018). These FSH inhibiting effects relied on the activation of the membrane estrogen receptor GPR30, and subsequent phosphorylation of protein kinase C (PKC) and MAPK proteins ERK and p38. One of the main mechanisms underlying DON toxicity involves the activation of MAPKs (Payros et al. 2016). Most studies have reported a rapid increase in phosphorylation of JNK, ERK and p38 proteins in various cell types following exposure to DON (Lee et al. 2019; Mishra et al. 2014; Zhang et al. 2020). However, the level of phosphorylated ERK decreased with increasing DON concentrations in piglet hippocampal nerve cells, indicating that the pattern of regulation of MAPKs may vary in a cell-type dependent manner (Wang et al. 2018).

GnRH is a central regulator of reproductive functions that acts by binding a G-protein coupled receptor on the surface of the gonadotrope cells. The GnRH binding transmits signals, mostly via the MAPK cascade, to increase the synthesis of LH and FSH (Melamed et al. 2012). Given the pivotal role of MAPKs for both the toxicity of DON and GnRH activation of the synthesis and release of gonadotropin by gonadotrope cells, we wanted to know, midstream in the HPG axis, what type of interference in GnRH-induced gonadotropin production could result from gonadotrope cells exposure to DON.

We observed herein a significant decrease of the viability of the gonadotrope-like cells even at the lowest tested concentration (1nM) of DON. Similar to the cell death mechanisms we previously reported in the intestinal mucosa, the cytotoxicity of DON toward the gonadotrope cells showed here appears to be driven by apoptosis (Payros et al. 2021). This cytotoxicity and the concentration ranges it occurred are relevant since *in vivo* experiments have established that the transport of DON across the blood-brain barrier is effective, with the toxin being detected in the brain within minutes of oral exposure (Behrens et al. 2015; Prelusky et al. 1990). Moreover, as the pituitary gland sits outside the blood-brain barrier, the tiny population of gonadotrope cells that constitute only 7% to 15% of the total number of anterior pituitary cells, are even exposed to higher concentrations of dietary DON compared to other parts of the brain. Considering the provisional maximum tolerable daily intake (PMTDI) of DON established by the Joint FAO/WHO Expert Committee on Food Additives, it is anticipated that the concentrations of DON in blood and cerebrospinal fluid will be 1.5 nM and 0.45 nM respectively (Maresca 2013). DON has been highlighted as of significant risk to human health, owing to the less than 30 safety factor between the doses of this mycotoxin susceptible to be present in the organism, in relation to its provisional maximum tolerable daily intake (PMTDI), and the doses affecting cell functions (Maresca and Fantini 2010; Muri et al. 2009). The 1 to 10 nM DON concentration range in this study showed clear cytotoxicity to LβT2 cells. Assuming that human gonadotrope cells have similar sensitivity to DON, the safety factor would be 0.7 to 20, positioning the pituitary as a prominent target for DON action.

We observed that exposure of gonadotrope-like cells to DON prior to stimulation by GnRH decreases the induced phosphorylation levels of Erk, but not p38. DON regulate MAPKs via the ribotoxic stress response, which involves the double-stranded RNA-activated protein kinase (PKR) and hemoitopoeitic cell kinase (Hck) as upstream transducers (Pestka 2008). Instead of PKR and Hck, GnRH activation of the phosphorylation of Erk and p38-MAPKs is mediated by several isoforms of PKC (Harris et al. 1997; Maccario et al. 2004). Moreover, the profile of the PKC isoforms involved in GnRH to p38MAPK phosphorylation differs from ERK1/2 activation. On top of the PKCβII, PKCδ and PKCε isoforms reported for GnRH activation of the phosphorylation of ERK in LβT2 cells, the GnRH induction of p38MAPK phosphorylation also involves PKCα (Mugami et al. 2018). Our results suggest that, contrary to PKCβII, PKCδ and PKCε, the activation of MAPKs by GnRH via PKCα may escape interference by DON. This hypothesis deserves to be investigated in the future.

The DON alteration of GnRH-induced phosphorylation of MAPKs in this study was followed by impaired induction of key regulatory genes for LH production (*Cga* and *Lhb*), and dose-dependent inhibition of the secretion of LH itself by gonadotrope cells. Although DON did not alter basal levels of *Fshb*, it would be inaccurate to conclude that DON has no effect on FSH secretion, since the experimental design of our study was specifically tailored to stimulate LH secretion rather than FSH (GnRH one pulse approach). Although produced by the same gonadotrope cell, the release of LH or FSH depends on specific GnRH pulsatility patterns, with LH release favored by fast GnRH pulse frequencies and FSH favored by slow pulse frequencies (Tsutsumi and Webster 2009). To allow simultaneous release of FSH and LH, mouse pituitary cells were pulsed for 2 minutes at a 10 nM GnRH peak pulse concentration with a 58-minute interpulse interval for 4 hours (Kim and Lawson 2015), which is drastically different from the 0.1 to 1 nM GnRH stimulation for 15 minutes or six hours used in the present study. Further investigations featuring the suitable pulsatility are needed to confirm if DON is also involved in the synthesis and release of FSH.

In conclusion, our study shows the first evidence that DON affects GnRH-induced signaling in gonadotrope-like cells through a mechanism that, at least in part, involves apoptosis and inhibition of GnRH-induced Erk phosphorylation. Most importantly, DON affected LH production by gonadotrope-like cells in a concentration-dependent manner. LH is a critical regulator of gonadal function in mammals and its altered synthesis or secretion can affect fertility in both sexes. This research broadens our knowledge of DON’s neurotoxic effects and brings a new depth to the potential neuroendocrine implications. This highlights the critical role of DON in compromising overall health, particularly its detrimental effects on the reproductive system, not only through direct effects on reproductive organs but also through interference with the control of critical hormones for reproduction. This provides novel insights and considerations for elucidating the toxicological mechanisms of DON.

## Acknowledgments

The authors thank Dr. Pamela Mellon (UC-San Diego) and Dr. Daniel J. Bernard (McGill University-Canada) for providing the LβT2 cells. The authors also thank the University of Virginia Ligand Core that provided assistance in LH measurement in cell culture supernatants. The University of Virginia Center for Research in Reproduction Ligand Assay and Analysis Core that is supported by the Eunice Kennedy Shriver NICHD/NIH Grant R24HD102061

## Funding

This work was supported by Foundation J.-Louis Lévesque. Lingchen Yang and Guodong Cai received a scholarship from China Scholarship Council.

## Financial interests

Authors declare they have no financial interests.

## Author Contributions

All authors contributed to the study conception and design. Conceptualization, Methodology and Funding acquisition were performed by Imourana Alassane-Kpembi; Formal analysis, Investigation, Data curation and Writing—original draft preparation were performed by Guodong Cai and Lingchen Yang; Writing—review and editing was performed by Francis Marien-Bourgeois, Derek Boerboom, Gustavo Zamberlam, and Imourana Alassane-Kpembi. Supervision was performed by Derek Boerboom, Gustavo Zamberlam and Imourana Alassane-Kpembi. All authors read and approved the final manuscript.

## Conflicts of Interest

The authors declare no conflict of interest.

## Notes

### Competing Interest Statement

The authors have declared no competing interest.

### Summary of Updates

Title revised, abstract revised, discussion section revised, figures revised and graphical abstract and figure captions added

